# Bacteria-phage (co)evolution is constrained in a synthetic community across multiple bacteria-phage pairs

**DOI:** 10.1101/2024.10.14.618343

**Authors:** Meaghan Castledine, Daniel Padfield, Marli Schoeman, Amy Berry, Angus Buckling

**Author notes:** **Corresponding author:** Meaghan Castledine.

## Abstract

Bacteriophages can be important drivers of bacterial densities, and therefore microbial community composition and function. These ecological interactions are likely to be greatly affected by evolutionary dynamics, because bacteria can rapidly evolve resistance to phage while phage can reciprocally evolve to increase infectivity. Most studies to date have explored eco-evolutionary dynamics using isolated pairs of bacteria-phage but in nature, multiple bacteria and phages coexist and (co)evolve simultaneously. How coevolution plays out in this context is poorly understood. Here, we examine how three coexisting soil bacteria (*Ochrobactrum* sp., *Pseudomonas* sp., and *Variovorax* sp.) interact and evolve with three species-specific bacteriophages over eight weeks of experimental evolution, both as host-parasite pairs in isolation and as a mixed community. Across all species phage resistance evolution was inhibited in polyculture, with the most pronounced effect on *Ochrobactrum*. Between bacteria-phage pairs there were also substantial differences in the effect of phage on host densities, and evolutionary dynamics including whether pairs coevolved. These contrasts emphasise the difficulty in generalising from monoculture to polyculture, and between bacteria-phage pairs to wider systems. Future studies should consider how multiple bacteria and phage pairs interact simultaneously to better understand how coevolutionary dynamics happen in natural communities.

**Importance:** This project is unique in examining evolutionary dynamics among coexisting bacteria and their phages, rather than focus on single focal species – this makes our work more applicable to natural contexts while still working with controlled synthetic communities. While it is commonly assumed that bacteria will evolve phage resistance and coevolve with phage, this may be uncommon in more complex communities due to reduced contact rates and/or reduced mutation rates. Furthermore, the contrast in population dynamics and ability to coevolve between bacteria-phage pairs highlights the need for more pairs to be studied. Over-reliance on model systems that are known to coevolve means we lack an understanding of how wider bacteria-phage pairs interact, and to what extent results can be generalised beyond these pairs.

## Introduction

Interactions between bacteria and phages can result in coevolution (1–3): the reciprocal evolution of bacterial resistance and phage infectivity. Coevolution can have profound impacts on bacteria-phage evolution and ecology, including driving population dynamics (4, 5) and the evolution of virulence of bacterial pathogens (6–8). While there is good evidence of coevolution in natural populations (9), bacteria-phage coevolution has been unequivocally demonstrated in several model systems including *Escherichia coli* (10), Pseudomonads (*P. fluorescens* (11)*, P. syringae* (12)), and *Staphylococcus aureus* (13).

Coevolutionary dynamics can be greatly affected by environmental conditions including the presence of competitors (14). While studies have characterised bacteria-phage coevolution with conspecifics (including in soil (14), plant leaves (15), digestive tracts (16, 17), and marine systems (18)), these studies typically focus on a single bacteria and phage pair (14, 17, 19–21), whereas in nature multiple bacteria-phage pairs coexist together.

Consequently, we do not understand how multiple bacteria and phages (co)evolve simultaneously. A general expectation is that pairwise reciprocal selection will be weakened in more complex communities. As densities are typically lower in communities, this will reduce encounter rates and therefore selection for resistance and infectivity (22, 23). Reduced densities and slower growth rates will reduce mutation supply rates thereby further slowing (co)evolution (1, 14, 24–26). Furthermore, competitors and parasites/predators can result in trade-offs in resistance to phage and maintaining competitive ability and/or defences to predators (20, 27, 28).

Here, we investigated how community context influences ecological and coevolutionary dynamics using a three-species bacterial community. This community consists of *Ochrobactrum* sp., *Pseudomonas* sp., and *Variovorax* sp., each of which has a species-specific lytic bacteriophage (29). We evolved these species for eight weeks in the presence and absence of other community members, with and without phage. Using resistance and time-shift assays (measuring bacteria-phage interactions both within and between time points), we examine how the presence of other bacteria species affected bacteria and phage coevolutionary interactions.

## Results

### Effects of phage are species-specific

With a three-species synthetic community, we sought to examine multiple bacteria-phage dynamics simultaneously in polyculture and monoculture. *Ochrobactrum* phage ORM_57 density was significantly greater in monoculture compared to polyculture (ANOVA comparing models with and without treatment: 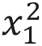 = 6.93, p = 0.008; Tukey HSD comparing densities in polyculture v monoculture: estimate = -2.2, t-ratio = -2.78, p = 0.019), with phages going below detectable densities in 5/6 replicates at week 6 and 4/6 replicates at week 8 (Figure 1a). On average phage densities declined from week two then stabilised (ANOVA comparing models with and without time: 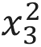 = 21.39, p < 0.001, Table S1; Figure 1a). Lower phage densities in polyculture may be attributable to *Ochrobactrum* densities being reduced under interspecific competition (Tukey HSD comparing *Ochrobactrum* density polyculture and monoculture (no phage) at week two: estimate = -0.444, t-ratio = -7.69, p < 0.001). Phages also significantly lowered bacterial densities in polyculture (ANOVA comparing models with and without treatment and phage interaction: 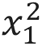 = 11.73, p < 0.001) while having no significant effect in monoculture (Tukey HSD: p > 0.05 for weeks two – six; Table S2; Figure 2a).

**Figure 1.**
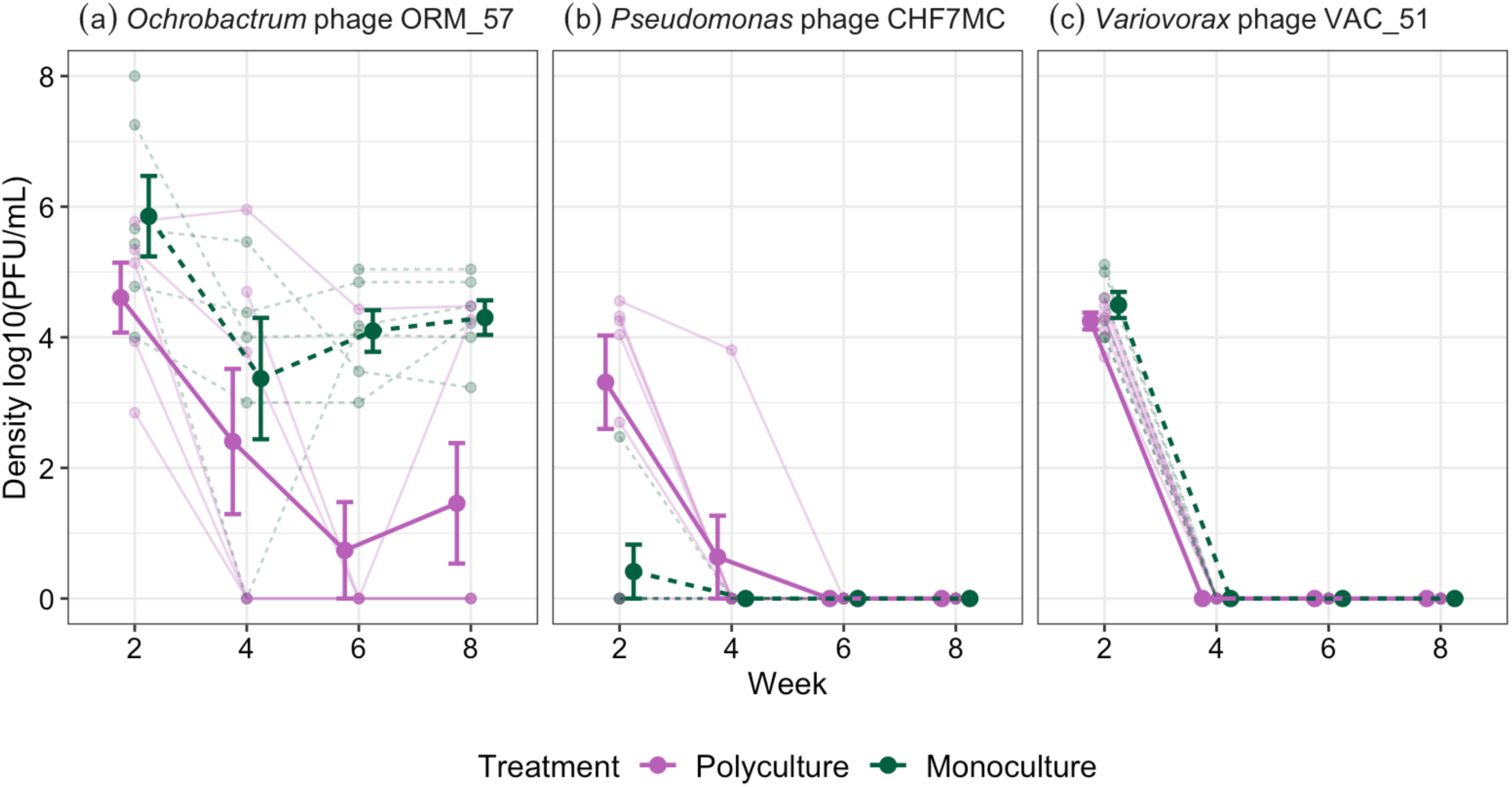
Phage density through time for (a) *Ochrobactrum* phage ORM_20, (b) *Pseudomonas* phage CHF7MC and (c) *Variovorax* phage VAC_51. Points with bars represent the means with standard errors. Small points represent separate treatment replicates. Lines connect points from the same treatment replicate.

**Figure 2.**
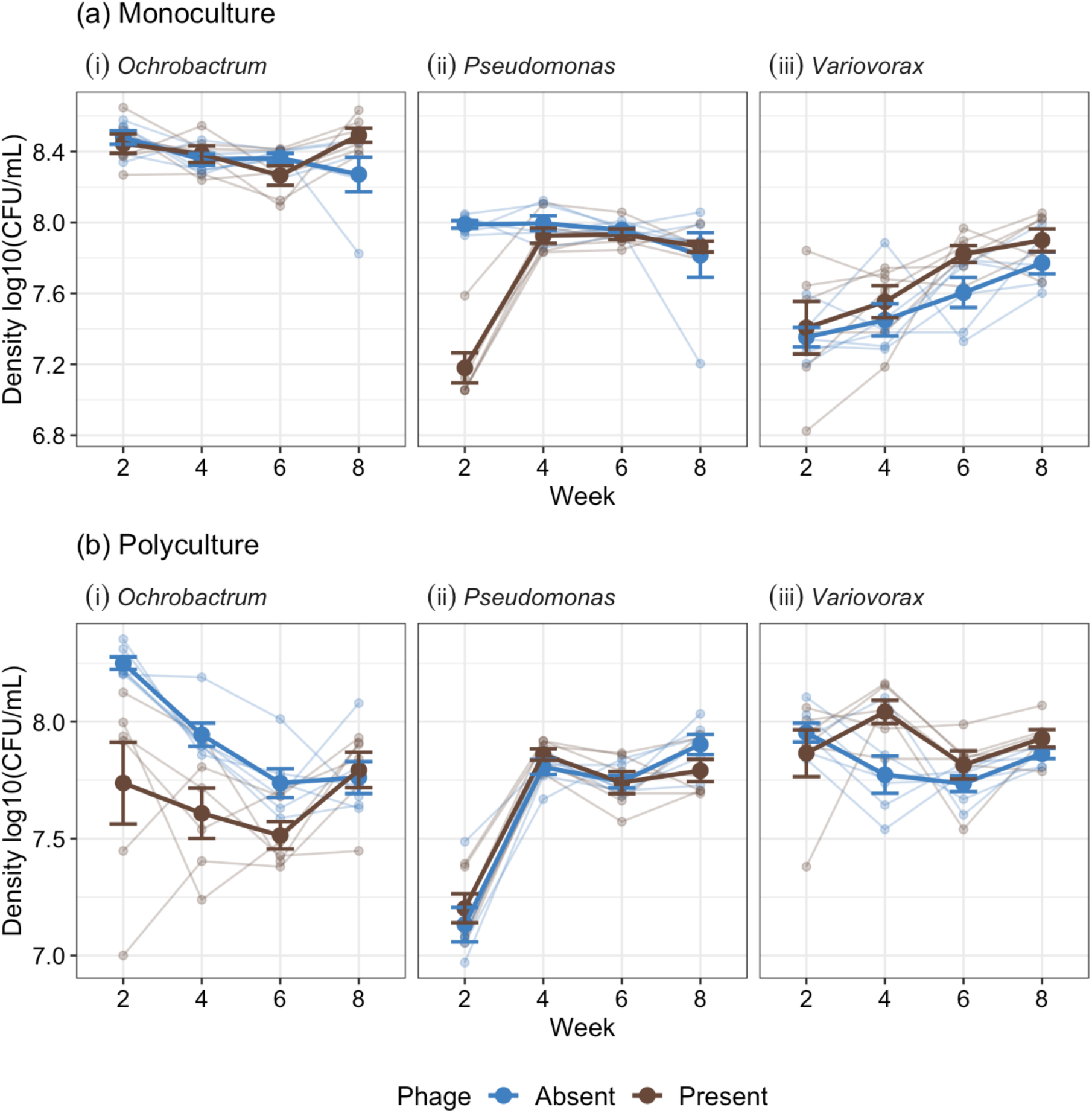
Changes in bacterial density through time with phages present or absent at the start of each experiment in (a) monoculture and (b) polyculture. Points with bars represent the means with standard errors. Small points represent separate treatment replicates. Lines connect points from the same treatment replicate.

For *Pseudomonas* and *Variovorax*, phages went extinct after 2-4 weeks (Figure 1b, c). Extinction of *Pseudomonas* phage CHF7MC was faster in monoculture compared to polyculture, with only one replicate containing phage in monoculture at week two compared to 5/6 replicates containing phage in polyculture (Figure 1b). The rapid extinction of *Pseudomonas* phage resulted in short-term effects of phage on bacterial densities in monoculture with bacteria densities recovering after week two (ANOVA comparing models with and without a three-way interaction between treatment, time and phage presence: 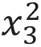 = 55.89, p < 0.001; Figure 2b, Table S3). In polyculture, phages did not significantly impact *Pseudomonas* density (Tukey HSD comparisons at week two between phage present/absent cultures: estimate = -0.07, t-ratio = -0.91, p-value = 0.801), however this may be due to interspecific competition providing bottom-up control of density (Tukey HSD comparisons at week two between phage present monocultures to phage absent polycultures: estimate = -0.05, t-ratio = -0.62, p = 0.926), indicating that effects of competition and phage were not additive.

*Variovorax* phage VAC_51 densities were not significantly affected by whether *Variovorax* was in mono-vs polyculture (ANOVA comparing models with and without treatment x time interaction: 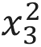 = 3.704, p = 0.295; independent effect of treatment: 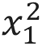 = 1.17, p = 0.279) and we detected no phage in any cultures at week four onwards (ANOVA comparing models with and without time: 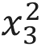 = 217.3, p < 0.001; Figure 1c). However, phages had a consistent positive effect on *Variovorax* densities, with cultures that had been exposed to phage reaching significantly higher densities than no-phage cultures (ANOVA comparing models with and without phage: 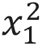 = 7.21, p = 0.007; Tukey HSD between no-phage and phage cultures: estimate = -0.103, t-ratio = -2.71, p-value = 0.013; Figure 2c). This effect was evident after phage extinction in both monoculture and polyculture (Figure 2c, Table S4).

Combined, we have observed species-specific effects of phage that also differed between monocultures and polycultures for two bacteria species. For *Pseudomonas*, densities were only affected by phage in monoculture and effects were absent once phages went extinct. *Ochrobactrum* was the only species to coexist with its phage, with bacteria and phage densities significantly lower in polyculture, while bacteria densities were unaffected by phage in monoculture. Comparatively, phage had a positive impact on *Variovorax* abundance in monoculture and polyculture, including after phage extinction.

### Phage resistance evolution is inhibited in polyculture

We next examined to what extent phage resistance was impacted by community context by comparing bacteria-phage evolution in monoculture and polyculture through time. As *Pseudomonas* and *Variovorax* phages went extinct after two-four weeks, we estimated phage resistance for all species at week two. Phage resistance levels were species specific (ANOVA comparing models with and without species x treatment interaction: F_2,29_ = 3.40, p = 0.047; Figure 3). For *Ochrobactrum*, phage resistance was inhibited in polyculture with no resistant clones while in monoculture 54.2% (SE (standard error) ±20.8) clones evolved resistance (Tukey HSD comparing resistance in polyculture to monoculture: estimate = -0.873, t-ratio = -3.983, p-value <0.001; Figure 3). Similarly, we found no phage resistance in *Pseudomonas* in polyculture while 6.94% (SE ±4.52) of clones were resistant in monoculture; although this was not significantly different (Tukey HSD: estimate = -0.157, t-ratio = -0.753, p-value = 0.457; Figure 3). *Variovorax* evolved 100% resistance in monoculture and 90.3% (SE ±6.6) resistance in polyculture which was non-significantly different (Tukey HSD: estimate = -0.215, t-ratio = -1.027, p-value = 0.313; Figure 3). These results show inhibited evolution of resistance across all species in polyculture, but this effect was largest for *Ochrobactrum*.

**Figure 3.**
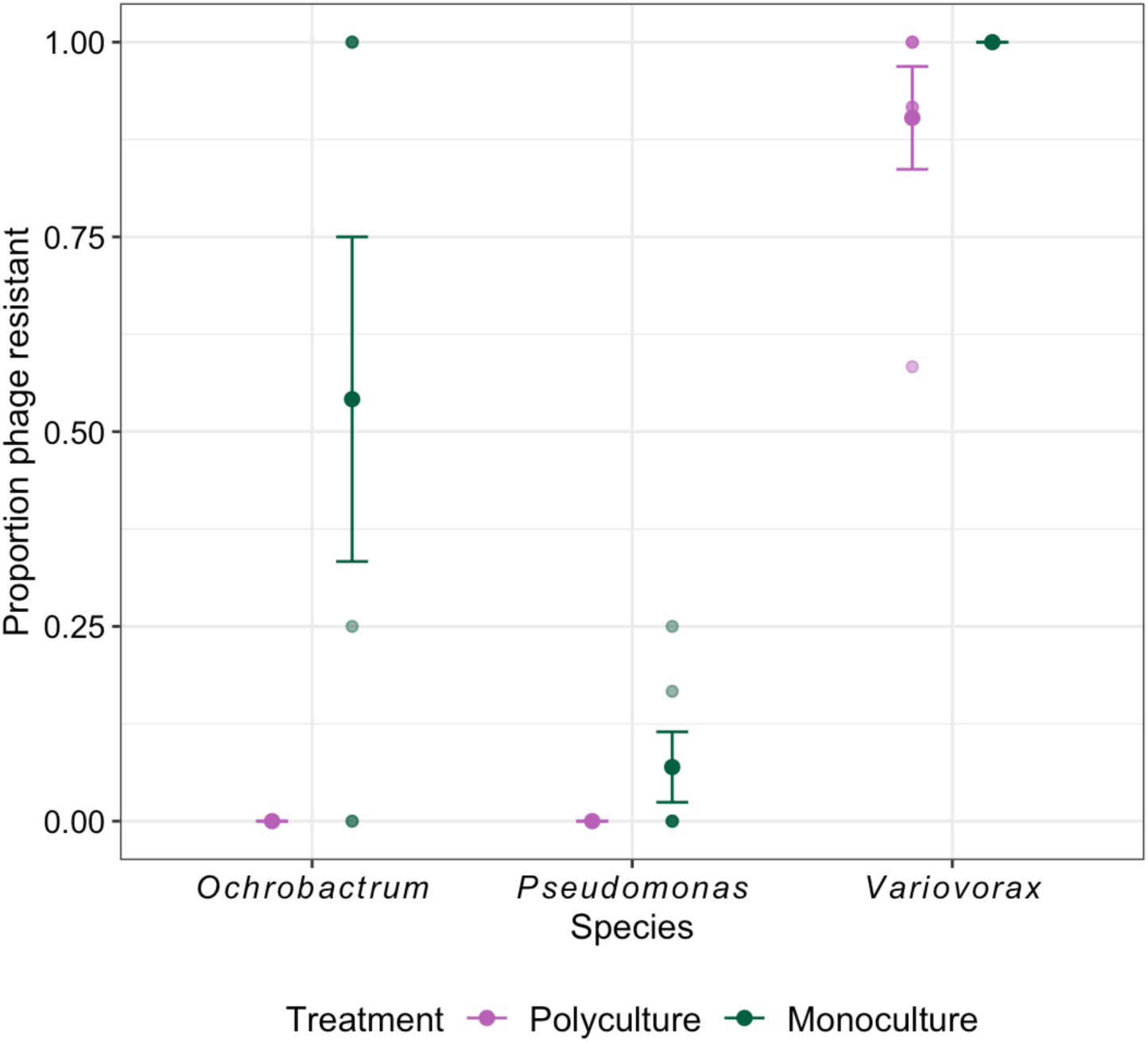
The proportion of phage resistant bacteria after two weeks evolving with phage in polyculture and monoculture. Points with bars represent the means with standard errors. Small points represent separate treatment replicates.

### Lack of resistance evolution in polyculture is not associated with resistance costs

It is not unusual for interspecific interactions to decrease phage resistance, with mechanisms ranging from reduced bacteria-phage contact rates, lower mutation supply rates or selection against less-fit resistant mutants. In *Ochrobactrum* especially, we found bacteria densities were significantly lower in polyculture when phage were added. With this result coinciding with lower rates of phage resistance, this could indicate context-specific resistance costs or density-effects reducing mutation supply rates and/or encounter rates (22, 26, 30). We examined whether interspecific competitors had competitively excluded phage resistant *Ochrobactrum* and *Variovorax* isolates owing to resistance costs to growth. This was not possible for *Pseudomonas* due to the low rates of resistance evolution. To this end, phage susceptible and resistant isolates were grown in monoculture and polyculture in the absence of phage and their growth rates compared. Here, the relative growth rate of was non-significantly different between ancestral, phage resistant and susceptible isolates for *Ochrobactrum* (ANOVA comparing models with and without treatment (monoculture, polyculture) x phage resistance: 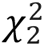 = 1.48, p = 0.476; phage resistance: 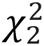 = 0.081, p = 0.994; Figure 4a) and *Variovorax* (ANOVA comparing models with and without treatment (monoculture, polyculture) x phage resistance: 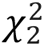 = 5.71, p = 0.057; phage resistance: 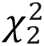 = 3.4, p = 0.183; Figure 4b). Both species growth rate was significantly lower in polyculture compared to monoculture (*Ochrobactrum*: ANOVA comparing models with and without treatment: 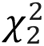 = 93.04, p < 0.001; Tukey HSD comparing growth rates in polyculture vs monoculture: estimate = -1.36, t-ratio = - 20.23, p < 0.001. *Variovorax*: ANOVA: 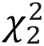 = 25.84, p < 0.001; Tukey HSD: estimate = -0.267, t-ratio = -10.63, p < 0.001). Consequently, lower rates of phage resistance in polyculture were not due to resistance costs and are more likely explained by lower mutation supply rates and/or lower encounter rates.

**Figure 4.**
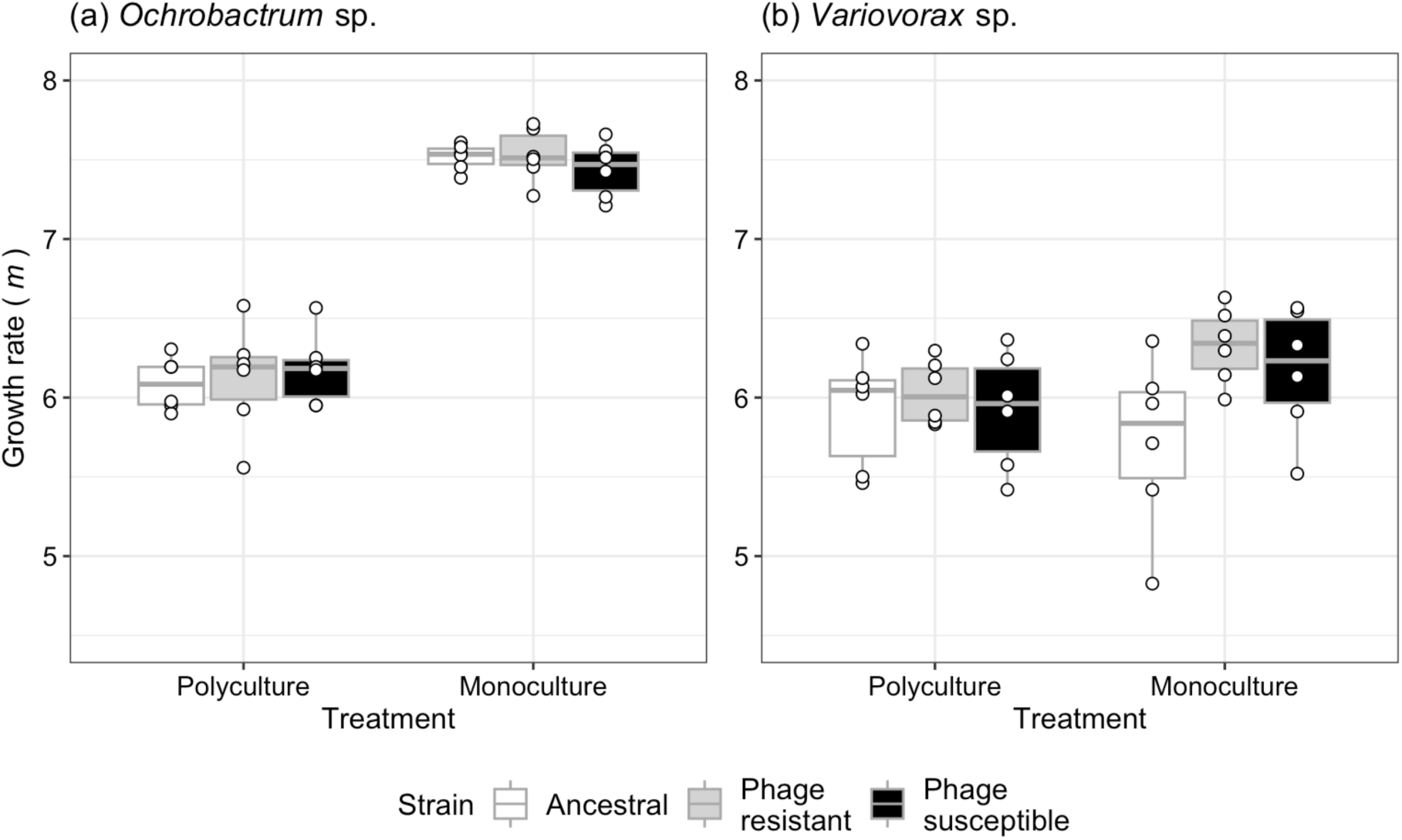
Growth rate of different (a) *Ochrobactrum* and (b) *Variovorax* strains, including ancestral, phage resistant and phage susceptible isolates, grown in polyculture and monoculture for one week. Points represent individual treatment replicates. Tops and bottoms of the bars represent the 75th and 25th percentiles of the data, white lines indicate the medians, and whiskers extend from their respective hinge to the smallest or largest value no further than 1.5× the interquartile range.

### Coevolutionary dynamics of *Ochrobactrum* and phage

As only *Ochrobactrum* coexisted with its phage (Figures 1a, 2a), we determined if coevolution had occurred by measuring the resistance of bacteria to phage both within and between time points. In polyculture, low frequency resistance (4.17%, SE ± 2.4%) was detected against ancestral phage only at week six, ruling out significant coevolution. In monoculture, 54.2% (SE ± 20.8%) of clones were resistant to the ancestral phage by week two. Phage infectivity increased (resistance decreased to evolved phage) through time (ANOVA comparing models with and without phage time: 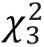 = 122.7, p <0.001; Tukey HSD comparing phage infectivity between ancestral phage, time-points two or four to six: p <0.001; Figure 5). Mean bacterial resistance declined at week four (ANOVA comparing models with and without bacterial time-point: 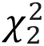 = 27.52, p < 0.001; Tukey HSD comparing resistance between weeks two and four: estimate = 1.12, z-ratio = 4.76, p <0.001; weeks four and six: estimate = - 0.972, z-ratio = -4.09, p <0.001) then increased at week six back to levels non-significantly different to week two (Tukey HSD weeks two and six: estimate = 0.15, z-ratio = 0.72, p = 0.752). However, bacteria from each time point showed similar patterns of resistance to phages isolated from past, contemporary, and future time-points (ANOVA comparing models with and without phage time x bacteria time interaction: 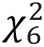 = 11.36, p = 0.078; Figure 5). These results suggest limited asymmetrical coevolution in monoculture with bacteria evolving resistance to phage (although resistance did not go to fixation), and phage slowly evolving increased infectivity against a subset of bacterial mutants through time.

**Figure 5.**
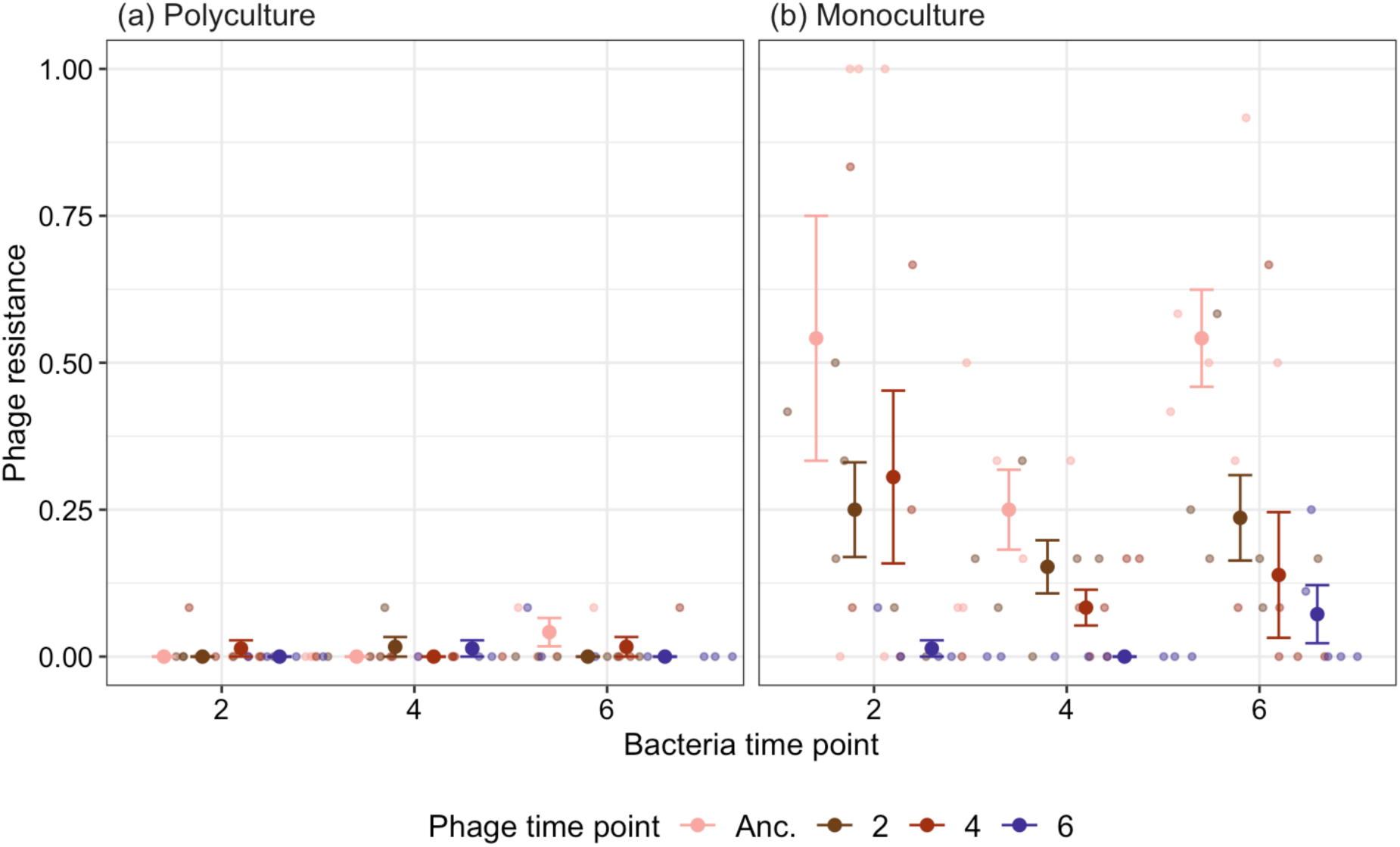
Changes in phage resistance (the proportion of phage resistant bacteria) in *Ochrobactrum* through time, demonstrated by assaying phages from different time-points against bacteria from different time-points. Points with bars represent the means with standard errors. Small points represent separate treatment replicates.

## Discussion

Over eight weeks of experimental evolution, we examined ecological and (co)evolutionary dynamics between three bacteria and phage pairs in monoculture (single pairs) and polyculture (all pairs coexisting). Overall, interactions between bacteria and phages were species-specific in effects on density, resistance evolution and (co)evolution. *Ochrobactrum* showed the biggest difference between monoculture and polyculture with densities only affected by phage in polyculture while coevolution only occurred in monoculture. Across all species, resistance evolution was inhibited in polyculture. While previous studies have examined bacteria and phage (co)evolution in polyculture (9), ours is unique in examining (co)evolutionary and ecological dynamics across all coexisting pairs as is the case in nature.

Ecological dynamics between bacteria and phage in polyculture and monoculture were species specific. While *Pseudomonas* densities’ were initially impacted by phage, densities recovered following phage extinction at week two. *Pseudomonas* phage CHF7MC is closely related to strain CHF7 which infects *P. syringae*, suggesting this phage may not be well-adapted to this *Pseudomonas* strain as extinction occurred without resistance (29). *Variovorax* densities significantly increased in the presence of phage, with this effect evident with fixation of phage resistance followed by phage extinction at week two. Phages have been shown to increase densities in other systems (14), but exploration of this mechanism is beyond the scope of this project. In contrast, *Ochrobactrum*’s densities were significantly reduced in polyculture when phages were present, and this was the only bacteria-phage pair to coexist for the study duration. This density decline is consistent with an interaction between phage lysis and interspecific competition as densities were unaffected in monoculture. Competitive interactions have been shown to increase effects of phage on bacteria density (31). Although *Ochrobactrum* evolved phage resistance in monoculture, which may have buffered densities against phage (32), its phage also increased in infectivity through time (92.8% week six isolates susceptible to contemporary phage) without affecting *Ochrobactrum* densities. As *Ochrobactrum* did not reciprocally increase in resistance, and densities did not decline, this would suggest that resistance alone did not buffer *Ochrobactrum* against the effects of its phage.

Across all species, resistance evolution was inhibited in polyculture. Consistent with previous research, phage resistance rates were lower in polyculture compared to monoculture (21). This effect was most evident for *Ochrobactrum* which coevolved with its phage in monoculture while not starting to evolve phage resistance until week six in polyculture (4.17%, SE ± 2.4%). *Variovorax’s* and *Ochrobactrum’s* lowered resistance in polyculture and was not attributed to phage resistance costs (in either polyculture or monoculture), and was most likely due to lowered bacteria-phage contact rates or mutation-supply rates (23, 33–35). Although we were unable to examine the mechanism of lowered resistance evolution in polyculture for *Pseudomonas*, bottom-up control of *Pseudomonas*’s density would have reduced contact and mutation rates in a similar manner to the other two species.

Bacteria-phage coevolution was evident for one pair, *Ochrobactrum* and its phage, in monoculture but not polyculture. While it is commonly assumed bacteria and phage coevolve, this is perhaps less likely in complex systems if resistance is unlikely to evolve ((9), although see (14, 15)). As studies typically focus on bacteria-phage pairs that are known to coevolve, it is less understood how common coevolution is in wider systems. Particularly in this case where phages are poorly infectious (*Pseudomonas*) or resistance can evolve rapidly and reach fixation (*Variovorax*), it is less likely that coevolution will occur.

That results contrast in monoculture and polyculture has important implications. Even in monoculture, each bacteria-phage interaction was species-specific and coevolution only occurred in one bacteria-phage pair. This emphasises the difficulty in generalising from model systems, and in focusing on model systems that behave predictably (e.g. always coevolve). Across all pairs, resistance evolution was inhibited in polyculture which further emphasises the difficulty in generalising results from less-natural monoculture conditions to complex communities. This is particularly important in the context of phage therapy where phages are screened in monoculture for ones which reduce bacteria densities the most, and result in the lowest rates of resistance (36). Although contact rates between bacteria and phage are typically high in phage therapy (therefore increasing selection), resistance may still be less likely *in vivo* owing to reduced bacteria replication rates and context dependent resistance costs (37). Although our community is still simplified compared to natural communities, by examining how multiple species evolve and interact simultaneously with their phages, our results have more ecological relevance than previous research focussing on single pairs.

## Materials and Methods

### Experimental evolution

Species isolates were originally obtained from soil and identified as: *Ochrobactrum* sp., *Pseudomonas* sp., and *Variovorax* sp. These species have unique colony morphologies when plated onto King’s medium agar (KB agar) and can stably coexist for several weeks (38, 39). In previous work, species-specific bacteriophages (*Ochrobactrum* phage ORM_20, *Pseudomonas* phage CHF7MC, *Variovorax* phage VAC_51) were identified and characterised for each species (29). The experimental treatments were: monoculture, monoculture with phage (each bacterial species individually with only its species-specific bacteriophage), polyculture (all three bacteria present) and polyculture with phage (all three bacteria with all three phages present). Each treatment was replicated six times. Each species was grown from one colony (clone) in isolation for two days in 6 mL growth media (1/64 Tryptic Soy Broth (TSB); diluted with demineralised H₂O), static, at 28 °C in 25 mL glass microcosms with loosened plastic lids. Species densities were normalised to ∼10^5^ CFU/μL as described previously (39). 10 μL of diluted culture (∼10^6^ CFUs) of each bacterial species were added to fresh vials. To phage-present cultures, ∼10^4^ PFUs (plaque forming units) were added (MOI 0.01). Serial 100-fold dilutions (60 μL culture into 6 mL growth media) took place every week for a total of eight weeks. Culture samples were cryogenically frozen at −70 °C in glycerol (final concentration: 25%) every transfer. Cultures were plated every 2^nd^ transfer onto KB agar and incubated for two days at 28 °C. Additionally, phage extractions were performed every 2nd transfer: 900 μL of culture was vortexed with 100 μL chloroform. Vials were then centrifuged at 14000 rpm (21100 *g*) for five minutes (mins) and the supernatant isolated. Phage densities were calculated via spot assays: phage cultures were diluted and 10 µL of each dilution was spotted onto soft agar overlays of each bacterium, with 100 µL of overnight bacterial culture added to 7 mL soft KB agar. If phages decreased below detectable densities, extracts were amplified in overnight cultures of ancestral strains of each species and re-spotted to confirm extinction.

### Measuring phage resistance and bacteria – phage coevolution

We estimated phage resistance to ancestral phage for each species from week two cultures. Twelve colonies were picked from each time point, for each bacterium within each treatment. Colonies were inoculated into 150 µL TSB and grown overnight at 28 °C in 96-well plates. Phage resistance was analysed using spot assays as above with ancestral phage.

As only *Ochrobactrum* and its phage coexisted for eight weeks across mono-and polyculture treatments, we were only able to characterise coevolution for this pair. Twelve bacterial colonies from each treatment replicate from weeks two, four and six were picked and grown as above. Where phage densities appeared extinct in week six replicates of polyculture lines, phages extracts were amplified overnight alongside ancestral *Ochrobactrum* to recover observable densities for coevolution analyses. Changes in phage resistance and infectivity were tested by spotting phages from different timepoints (ancestral, two, four and six) against bacteria from the same, past or future timepoints. If bacteria and phage are coevolving via arms-race dynamics, we expect bacteria to be more resistant to phage from the past and more susceptible to phage from future timepoints (11, 40). If dynamics follow a fluctuating-selection dynamic, we expect bacteria to have greater resistance to contemporary phage populations (1).

### Cost of resistance assay

Next, we considered if resistance to phage was costly to *Ochrobactrum’*s and *Variovorax’s* growth rate in monoculture and polyculture. This was not possible for *Pseudomonas* as phage resistance only emerged in two replicates, therefore not providing enough clones from independent replicates. Phage resistant, phage susceptible and ancestral *Ochrobactrum* and *Variovorax* isolates were grown in monoculture and polyculture with ancestral strains of the other two species, in the absence of phage (six replicates per treatment). For *Ochrobactrum*, six phage resistant and six phage susceptible isolates were isolated from independent replicates (1 resistant and 1 susceptible clone from each replicate) from week two monoculture with phage lines where both genotypes coexisted. For *Variovorax*, six independent phage resistant clones and six independent phage susceptible clones were isolated from phage and no phage week two monoculture with phage evolution lines respectively. ‘No phage’ lines were selected for isolation of phage susceptible clones as phage resistance was too high in ‘phage’ lines for enough individual susceptible clones to be isolated. Ancestral, phage resistant and phage susceptible isolates were individually grown for two days at 28 °C, shaking (180 r.p.m.) in 6 mL 1/64 TSB. Ancestral *Ochrobactrum*, *Pseudomonas* and *Variovorax* isolates were also grown in the same conditions. Isolate densities were normalised to 10^5^ CFUs/μL and 10 μL added to relevant vials. Cultures were grown static for 1 week at 28 °C, then frozen as previous and plated from frozen. We calculated growth rate as ln(N_1_/ N_0_)/ *t* where N_1_ is the final density, N_0_ is starting density and *t* is the time given for growth. As *t* is equal across experiments (1 week), this equation simplifies to *m* = ln(N_1_/ N_0_). This measure equates to an estimate of the Malthusian parameter if population were growing exponentially, but they were unlikely to be for the duration of the week-long assay. 1 week is selected as appropriate for comparing results to experimental evolution where cultures are transferred weekly.

### Statistical analyses

All data were analysed using R (v.4.2.1) in RStudio (41) and all plots were made using the package ‘*ggplot2*’ (42). Model simplification was conducted using likelihood ratio tests and Tukey’s post hoc multiple comparison tests were done using the R package ‘*emmeans*’ (43). In a linear mixed effects model, bacterial density (log10 CFU/mL) is analysed against interacting fixed effects of phage presence, treatment (monoculture/polyculture) and time with a random effect of treatment replicate. Similarly, separate models were run for each phage strain analysing phage density (log10 PFU/mL) against interacting fixed effects of treatment and time, with a random effect of treatment replicate. As we were not interested in comparing densities between species, but rather the effects of phage on individual species between treatments, individual models were run for each species.

Phage resistance was analysed for week two cultures in a linear model with the proportion of phage resistant isolates (arcsine transformed) analysed against interacting effects of treatment (monoculture / polyculture) and species. To assess whether *Ochrobactrum* had coevolved with phage, the proportion of phage susceptible isolates was analysed in a generalised linear model mixed effects model with interacting fixed effects of phage time, bacteria time, and treatment (monoculture or polyculture) with a binomial error structure and a random effect of treatment replicate.

The effect of phage resistance on *Ochrobactrum* and *Variovorax* growth rate (*m*) was estimate using separate linear mixed effects models for each species. Growth rate (*m*) was analysed against interacting fixed effects of treatment (monoculture, polyculture) and phage resistance (ancestral, resistant, susceptible). For *Ochrobactrum*, a random effect of clonal identity (same clones used across different treatments) nested within the treatment replicate clones were isolated from was included. For *Variovorax*, a random effect of clonal identity was included.

## Author contributions

MC and AnB conceived and designed the study. Experiments conducted by MC, MS and AmB. Data analysis conducted by MC. All authors contributed to the writing of the manuscript.

## Acknowledgements

We thank Stineke van Houte and Michael Brockhurst for helpful feedback on a manuscript draft. This work was funded by grant no. MR/N0137941/1 for the Great West 4 BIOMED Medical Research Council Doctoral Training Partnership, awarded to the Universities of Bath, Bristol, Cardiff, and Exeter from the Medical Research Council/UK Research and Innovation, awarded to MC. This work was supported by NERC awards NE/V012347/1 and NE/S000771/1 awarded to AB. DP is funded by a NERC independent research fellowship (NE/W008890/1).

## Data availability

R code and data are deposited on GitHub (github.com/mcastledine96/Polyculture_suppresses_coevolution_2024)

